# Differentiated neuroblastoma cells remain epigenetically poised for de-differentiation to an immature state

**DOI:** 10.1101/2022.07.10.499470

**Authors:** Richard A. Guyer, Nicole Picard, Jessica L. Mueller, Andrew J. Murphy, Kristine M. Cornejo, Ryo Hotta, Allan M. Goldstein

**Author notes:** Corresponding author: Allan M. Goldstein, MD. These authors contributed equally to this work and share first authorship (Richard A. Guyer, Nicole Picard).

## Abstract

Neuroblastoma is the most common extracranial solid tumor of childhood and accounts for a significant share of childhood cancer deaths. Prior studies utilizing RNA sequencing of bulk tumor populations showed two predominant cell states characterized by high and low expression of neuronal genes. Although cells respond to treatment by altering their gene expression, it is unclear whether this reflects shifting balances of distinct subpopulations or plasticity of individual cells. Using neuroblastoma cell lines lacking *MYCN* amplification, we show that the antigen CD49b distinguishes these subpopulations. CD49b expression marks proliferative cells with an immature gene expression program, while CD49b-negative cells express differentiated neuronal marker genes and are quiescent. Sorted populations spontaneously switch between CD49b expression states in culture, and CD49b-negative cells can generate rapidly growing, CD49b-positive tumors in mice. We profiled H3K27ac to identify enhancers and super enhancers that are specifically active in each population and find that CD49b-negative cells maintain the priming H3K4me1 mark at elements that are active in CD49b-high cells. Improper maintenance of primed enhancer elements thus may underlie cellular plasticity in neuroblastoma, representing potential therapeutic targets for this lethal tumor.

**Summary Statement:** This study demonstrates that neuroblastoma cells can interconvert between a state characterized by expression of neuronal genes and a de-differentiated state.

## Introduction

Neuroblastoma is the most common extracranial solid tumor in children, and accounts for 15% of pediatric cancer deaths annually (Newman et al., 2019). These tumors arise when normal differentiation of neural crest cells into sympathetic neurons of the peripheral nervous system is disrupted (Kildisiute et al., 2021). The disease is stratified based on clinical and molecular characteristics, with high-risk tumors carrying a dismal prognosis (Cohn et al., 2016). Although *MYCN* amplification is the most common mutation found in high-risk lesions, over half of these tumors do not display *MYCN* amplification (Lee et al., 2018; Yanishevski et al., 2020).

Cellular identity is defined by the proteins that have been translated at any moment, and protein translation requires transcription of genomic information into RNA. RNA expression is determined by the enhancer elements a cell has selected (Long et al., 2016). Primed enhancers, marked by mono-or dimethylation of lysine 4 on histone 3 (H3K4me1/2), are not actively engaged in promoting transcription, while active enhancers denoted by the addition of acetylation on lysine 27 on histone 3 (H3K27ac) are bound by transcription factors and replication machinery (Creyghton et al., 2010). Stem and progenitor cells are characterized by a broad repertoire of primed enhancers that can be activated to trigger a change in transcriptional state (Crispatzu et al., 2021; Rada-Iglesias et al., 2011). As cells progress through differentiation options, enhancers are decommissioned via loss of H3K4me1/2 marks to limit fate potential (Tao et al., 2021; Whyte et al., 2012). Dysregulation of epigenetic pathways, including aberrant enhancer activity, is common in cancer (Helmsauer et al., 2020; Okabe and Kaneda, 2021).

Super enhancers (SEs) are large regions of chromatin that are densely bound by transcription factors and are strongly marked by H3K27ac (Parker et al., 2013; Whyte et al., 2013). Genes controlled by super enhancers are highly expressed, and often sit at the apex of networks that establish cell identity (Whyte et al., 2013). Cancer cells, including neuroblastoma cells, are particularly sensitive to altered transcription of SE-controlled genes (Lovén et al., 2013). Identification of SEs is thus valuable for determining target points to disrupt tumors and understanding how SEs are dysregulated may provide insights regarding mechanisms of tumor initiation and progression.

There are two predominant biological states of neuroblastoma cells: an undifferentiated mesenchymal state and an adrenergic state more closely resembling differentiated, committed sympathetic neurons (Gartlgruber et al., 2021; van Groningen et al., 2017). Gene signatures associated with these states have prognostic value, with the mesenchymal phenotype being associated with worse outcomes, and relapsed lesions also being enriched for markers of the mesenchymal state (Gartlgruber et al., 2021; van Groningen et al., 2017). Tumor cell lines treated with radiation or chemotherapeutic agents have been shown to adopt a mesenchymal gene expression profile, a process which involves activation of NOTCH signaling (Boeva et al., 2017; van Groningen et al., 2019). This seemingly conflicts with longstanding evidence that post-treatment tumors often undergo histologic maturation (Finklestein et al., 1979; Grosfeld et al., 1978). However, despite histologic maturation with therapy, many patients relapse and succumb to their disease, raising the possibility that mature cells seen soon after therapy are replaced by cells with less differentiated features. Moreover, it is unclear whether neuroblastoma plasticity represents clonal evolution with shifting balances among distinct tumor subpopulations or plasticity of individual cells.

The present study was undertaken to determine whether neuroblastoma cells lacking *MYCN* amplification include heterogeneous cellular phenotypes. We investigated whether the cell-surface marker CD49b (integrin alpha 2, *Itga2*), which marks proliferative neural crest progenitor cells (Abe et al., 2016; Joseph et al., 2011), distinguishes neuroblastoma populations with distinct gene expression programs. We found that cells lacking CD49b expression are quiescent, express RNAs encoding adrenergic transcription factors, and transcribe neuronal marker genes. In contrast, CD49b-expressing cells are proliferative, transcribe transcription factor genes associated with the mesenchymal cell state, and do not express neuronal genes. As expected, different complements of active enhancers and SEs are associated with these populations. Intriguingly, we found that CD49b-low cells, which otherwise show many hallmarks of mature neurons, maintain the priming H3K4me1 mark at many enhancers and SEs that are active only in CD49b-high cells, suggesting that mature cells retain an abnormal ability to de-differentiate. Importantly, cells with either phenotype can give rise to the opposite cell type in culture. These results suggest that a bidirectional differentiation hierarchy exists in neuroblastoma, likely due to failure to decommission enhancer and SE elements. Defining the epigenetic mechanisms that restrict neural crest cell fate may thus be critical for understanding and treating this aggressive childhood disease.

## Results

### Neuroblastoma Cells Express the integrin CD49b

Because CD49b marks neuronal progenitor cells in neural crest-derived lineages but is not expressed on differentiated neurons (Belkind-Gerson et al., 2015; Joseph et al., 2011; Morarach et al., 2021), we hypothesized that this antigen would distinguish immature and mature neuroblastoma cells. We first assessed expression of the *Itga2* gene, which encodes for the CD49b antigen, in mouse 3T3 Swiss (3T3) fibroblast cells, as well as in the murine neuroblastoma cell line Neuro-2a (N2a), which lacks *Mycn* amplification. As a positive control, we also assessed for expression of *Ngfr* and *Phox2b*, two genes known to be highly expressed in neuroblastoma (Baker et al., 1989; Boeva et al., 2017).

Consistent with our hypothesis, N2a cells display markedly higher transcript levels of *Itga2* than do 3T3 cells (Fig. 1A). We followed this result by assessing cell surface expression of CD49b on N2a cells, as well as on human SH-SY5Y cells, a patient-derived neuroblastoma cell line that also lacks *MYCN* amplification. Both lines are heterogeneous with respect to CD49b cell-surface expression (Fig. 1B,C). We noted that N2a cells have a continuum of expression, from cells lacking the antigen to those with high expression, while SH-SY5Y cells have more clearly distinct CD49b-negative and CD49b-positive populations.

**Figure 1.**
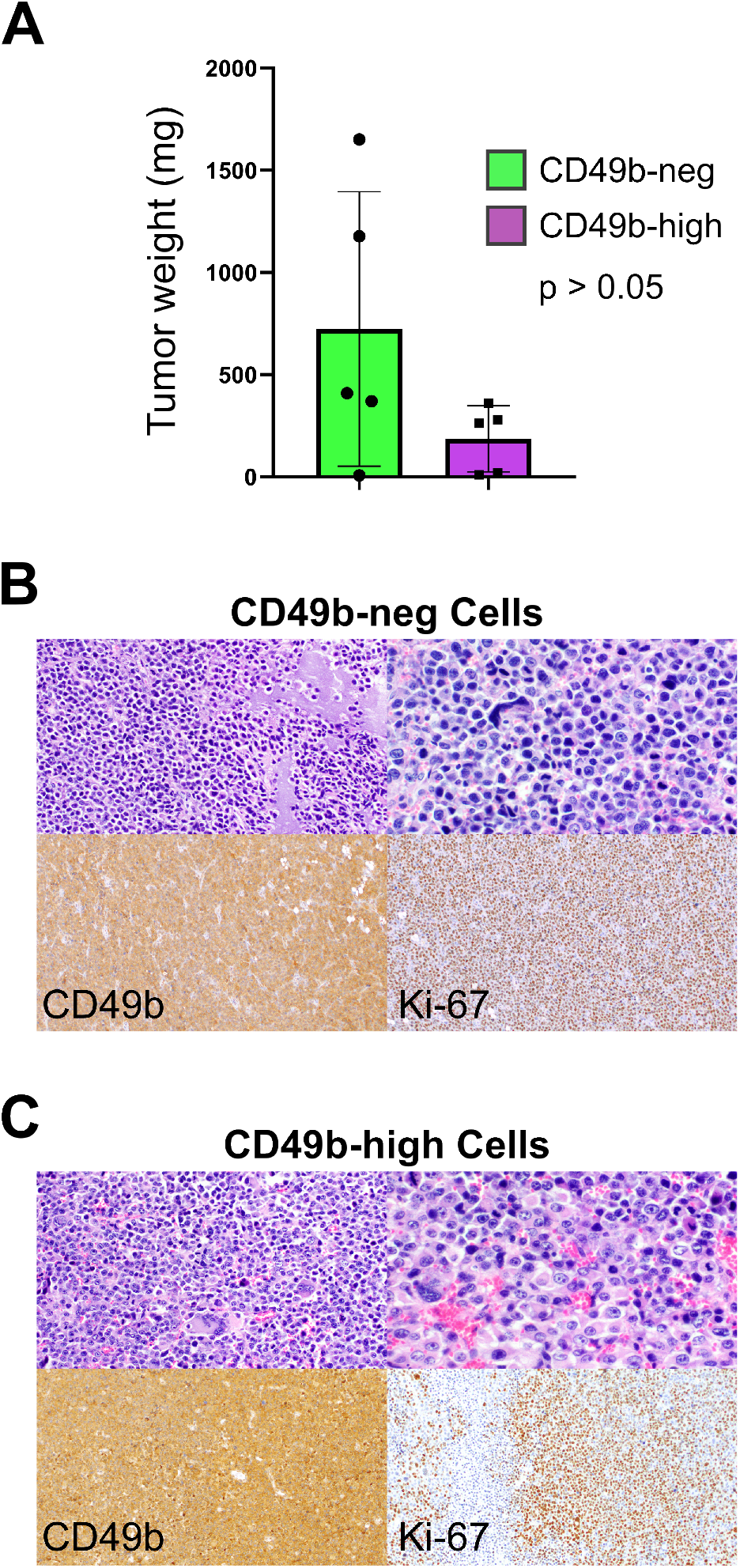
Neuroblastoma cells express the CD49b cell-surface antigen. A) qPCR for the indicated genes in the mouse Neuro-2a neuroblastoma cell line, as well as the mouse 3T3 Swiss fibroblast cell line. Bars indicate mean, dots indicate individual replicates, and error bars indicate standard deviation. *Gapdh* expression was used for normalization. Two-tailed t-test used to calculate p-value. B) Representative flow cytometry plots showing CD49b antigen expression on mouse Neuro-2a neuroblastoma cells. C) Representative flow cytometry plots showing CD49b antigen expression on human SH-SY5H neuroblastoma cells.

### CD49b Expression Distinguishes Neuroblastoma Cell States

To determine whether variation in CD49b expression reflects the differentiation state of neuroblastoma cells, we sorted both N2a and SH-SY5Y cells based on CD49b expression. Because CD49b does not demarcate discrete populations of N2a cells, we sorted these cells into the lowest and highest expression quartiles based on staining for the antigen (Fig. S1A). We henceforth called these populations CD49b-neg and CD49-high, respectively, since the lowest 25% of N2a cells based on CD49b staining displayed similar signal as the unstained control (Fig S1A). The intermediate 50% of N2a cells, which express low levels of the antigen, are referred to as CD49b-low. In contrast, SH-SY5Y cells were sorted into distinct populations with and without detectable CD49b antigen, which we respectively call C49b-pos and CD49b-neg (Fig. S1B).

After sorting, we isolated RNA and performed qPCR for selected neuronal marker genes (Fig. 2A,B). Consistent with CD49b’s role as a marker of immature neuronal precursors in other neural crest-derived lineages, we found diminished expression of the neuronal genes *Elavl4, Phox2a, Phox2b, Snap25*, and *Actl6b*, although only *Elavl4* and *Phox2b* reached statistical significance (p < 0.05) in N2a cells (Fig. 2A). In SH-SY5Y cells, transcripts of the human genes *ELAVL4, PHOX2A, PHOX2B, TUBB3, SNAP25*, and *ACTL6B* were diminished, with all except *PHOX2A* reaching statistical significance (Fig. 2B).

**Figure 2.**
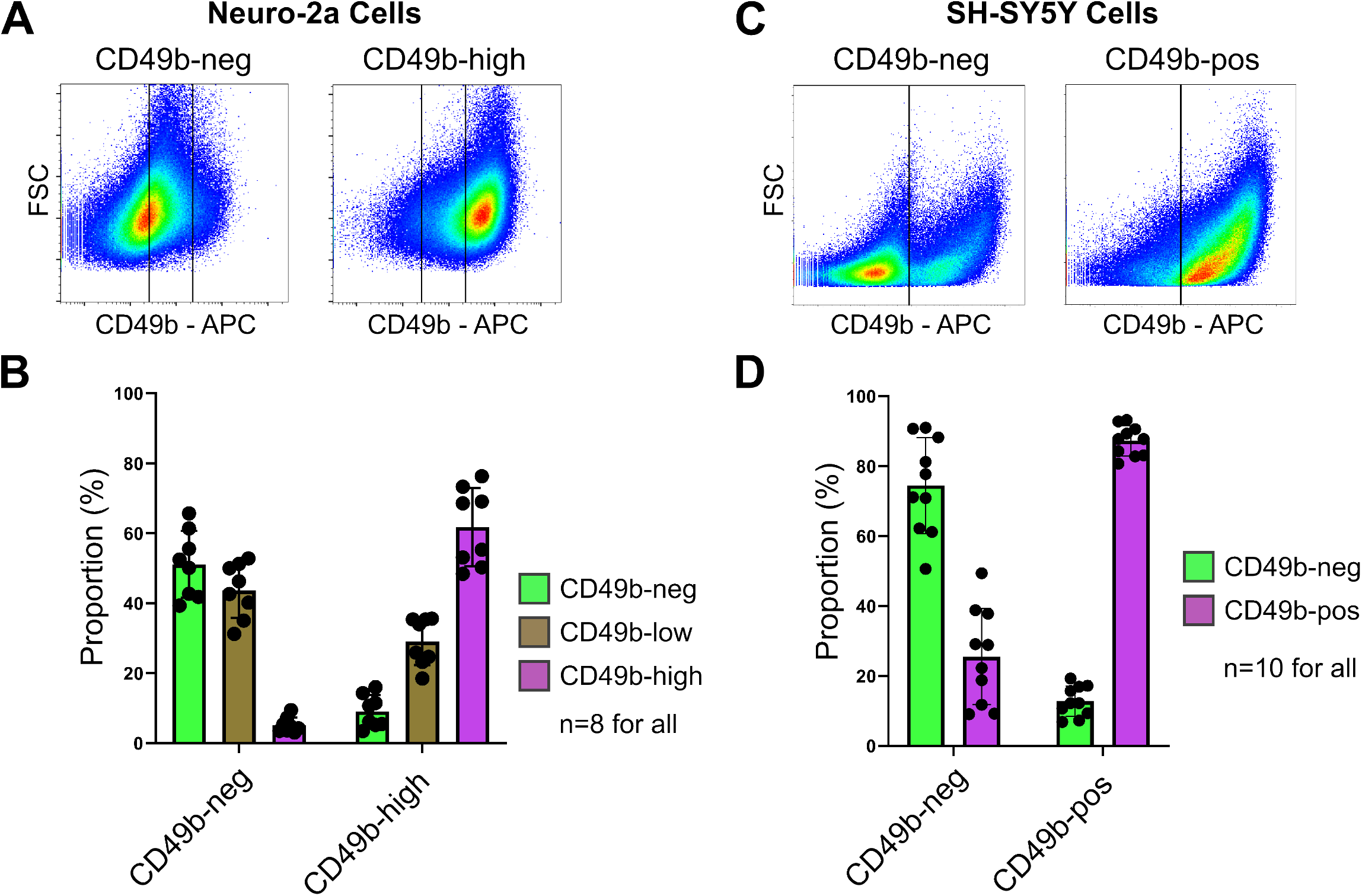
CD49b expression distinguishes transcriptionally disparate subpopulations within neuroblastoma cell lines. A) qPCR for the indicated genes in N2a cells sorted into CD49b-neg and CD49b-high fractions. Bars indicate mean, dots indicate individual replicates, and error bars indicate standard deviation. *Gapdh* expression was used for normalization. Two-tailed t-tests used to calculate p-value. B) qPCR for the indicated genes in SH-SY5Y cells sorted into CD49b-neg and CD49b-pos fractions. Bars indicate mean, dots indicate individual replicates, and error bars indicate standard deviation. *GAPDH* expression was used for normalization. Two-tailed t-tests used to calculate p-value. C) Volcano plot showing differentially expressed genes in CD49b-neg N2a cells relative to CD49b-high N2a cells identified by poly(A)-enriched RNA sequencing. 4251 genes had and absolute value log2-fold expression difference > 1 and p-value < 0.001. D) Volcano plot showing differentially expressed genes in CD49b-neg SH-SY5Y cells relative to CD49b-pos SH-SY5Y cells identified by poly(A)-enriched RNA sequencing. 8409 genes had and absolute value log2-fold expression difference > 1 and p-value < 0.001. E) Heatmap showing enrichment of selected neuronal marker genes in CD49b-high N2a cells relative to CD49b-neg N2a cells. For E and F, Heatmaps reflect z-score of log2-scale differences in gene expression. F) Heatmap showing enrichment of selected neuronal marker genes in CD49b-pos SH-SY5Y cells relative to CD49b-neg SH-SY5Y cells.

We next performed poly(A)-enriched RNA sequencing on sorted CD49b populations in both cell lines. These experiments were undertaken to test, in an unbiased manner, whether CD49b marks biologically distinct neuroblastoma cells. We found striking differences, with 4251 genes in N2a cells and 8409 genes in SH-SY5Y cells achieving the predetermined thresholds of greater than 2-fold difference in gene expression and p<0.0001 (Fig. 2C,D). To query whether this reflects neuronal differentiation status, we compared our RNA sequencing replicates based on expression of selected neuronal marker genes. Consistent with our qPCR data, CD49b-neg cells in both the N2a and SH-SY5Y lines were markedly enriched for neuronal genes (Fig. 2E,F). We also examined genes encoding transcription factors that have been associated with the adrenergic and mesenchymal cell states in the literature (van Groningen et al., 2017). The adrenergic factors, which include many genes associated with neurogenesis in the neural crest, were enriched in the CD49b-neg population in both cell lines (Fig.S2A). In contrast, the mesenchymal factors displayed notably higher transcript levels in the CD49b-high population in N2a cells and CD49b-pos population in SH-SY5Y cells (Fig. S2B). These data confirm that CD49b distinguishes distinct populations among neuroblastoma cells, and that lack of CD49b indicates cells with a transcriptional program characteristic of differentiated neurons.

### Distinct Signaling Pathways Characterize Cells Distinguished by CD49b Expression

To further validate that CD49b expression identifies distinct cell groups, we performed Gene Set Enrichment Analysis (GSEA) to identify biological pathways that are preferentially active in one population or the other. The results, shown in Tables S2 and S3, were remarkably similar for both cell lines. We focused on the PI3K-Akt and Cytokine-Cytokine Receptor Interaction pathways, both of which had significant differences in both cell lines and which are known to impact neuroblastoma cell phenotypes (Cotterman and Knoepfler, 2009; Hatzi et al., 2002; Paul et al., 2013). Enrichment plots and heatmaps of genes associated with these pathways show significant enrichment in the CD49b-high and CD49b-pos populations in N2a and SH-SY5Y cells, respectively (Fig. S3A-D). We used flow cytometry to confirm that CD49b-positive cells have higher levels of active, phosphorylated Akt/AKT in each cell line, as well as a higher percentage of cells that express the gp130 cytokine receptor (Fig. S3E,F). These results provide additional evidence that CD49b expression distinguishes neuroblastoma cells with distinct biological traits.

### Clinical Neuroblastoma Specimens Encompass Similar Heterogeneity as Cell Lines

Experiments with tumor cell lines may not reflect the biology of clinical disease. To assess the clinical relevance of our results, we assessed single-cell gene expression in 34,501 tumor cells isolated from five children with high-risk neuroblastomas without *MYCN* amplification. Hierarchical clustering demonstrates that these tumors contain a heterogenous mix of cells, with a subset enriched for common neuronal marker genes (Fig. 3A). We inferred cell cycle status of these genes based on expression of cell cycle-associated genes, as previously described (Tirosh et al., 2016). Strikingly, except for *PHOX2B*, all the neuronal marker genes we assessed are enriched in quiescent cells, with diminished transcript levels among cells in the S, G2, and M phases of the cell cycle (Fig. 3B). We confirmed this result by sorting the human SH-SY5Y cell line into CD49b-pos and CD49b-neg populations, plating equal numbers of cells from each group, and assessing proliferation via a luminescence assay 2, 5, and 7 days later. The CD49b-pos population showed significantly greater proliferation at all time points (p<0.005; Fig. 3C). This demonstrates that primary neuroblastomas have similar transcriptional heterogeneity as the N2a and SH-SY5Y cell lines, and that cells with a neuronal gene expression signature are quiescent.

**Figure 3.**
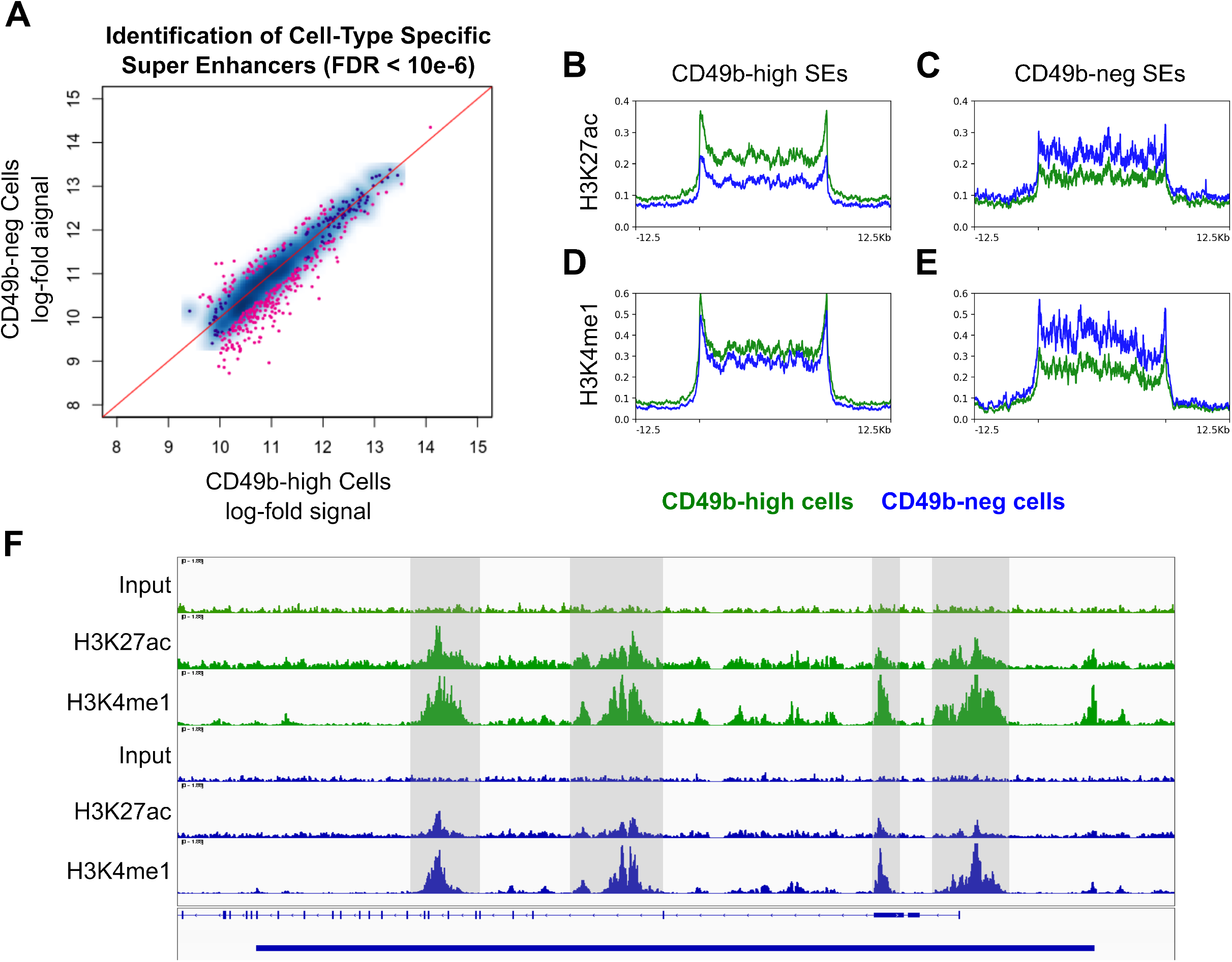
Single-cell RNA sequencing confirms primary neuroblastoma lesions are heterogeneous and expression of neuronal genes correlates with cell cycle status. A) Single-cell heatmap showing expression of the indicated neuronal markers in 34,501 tumor cells isolated from 5 children with high-risk tumors without *MYCN* amplification. B) Dot plot showing the proportion of cells expressing the indicated neuronal marker genes, stratified by cell cycle status inferred from gene expression patterns. Dot size indicates the proportion of cells expressing each gene, and color indicates the relative level of expression. C) Proliferation in culture of SH-SY5Y cells sorted into CD49b-pos and CD49b-neg fractions. Two-tailed t-tests used to calculate p-value.

### CD49b-neg N2a cells maintain H3K4me1 priming marks at enhancers active in CD49b-high cells

Gene expression is determined by the enhancer elements active in cells. Given the striking differences in gene expression among subpopulations of neuroblastoma cells, we hypothesized that cells with different levels of CD49b antigen expression would have distinct enhancer profiles. Because acquisition of a neuronal gene signature, including *ACTL6B* (*Actl6b* in mice), is associated with terminal differentiation during normal neurogenesis (Morarach et al., 2021; Yoo et al., 2017), we also hypothesized that CD49b-neg cells would decommission enhancers that are active in CD49b-pos/high cells. We used CUT&RUN to globally assay H3K27ac and H3K4me1 in CD49b-neg and CD49b-high populations of N2a cells. We first identified enhancers active in each population by assessing H3K27ac signal. Using a false discovery rate (FDR) of <0.001, we identified 3622 enhancers specifically active in one population or the other, including 2225 enhancers active in CD49b-high cells and 1397 enhancers active in CD49b-neg cells (Fig. 4A). As anticipated, enhancer regions activated in each population displayed markedly diminished H3K27ac signal in the other population (Fig. 4B,D).

**Figure 4.**
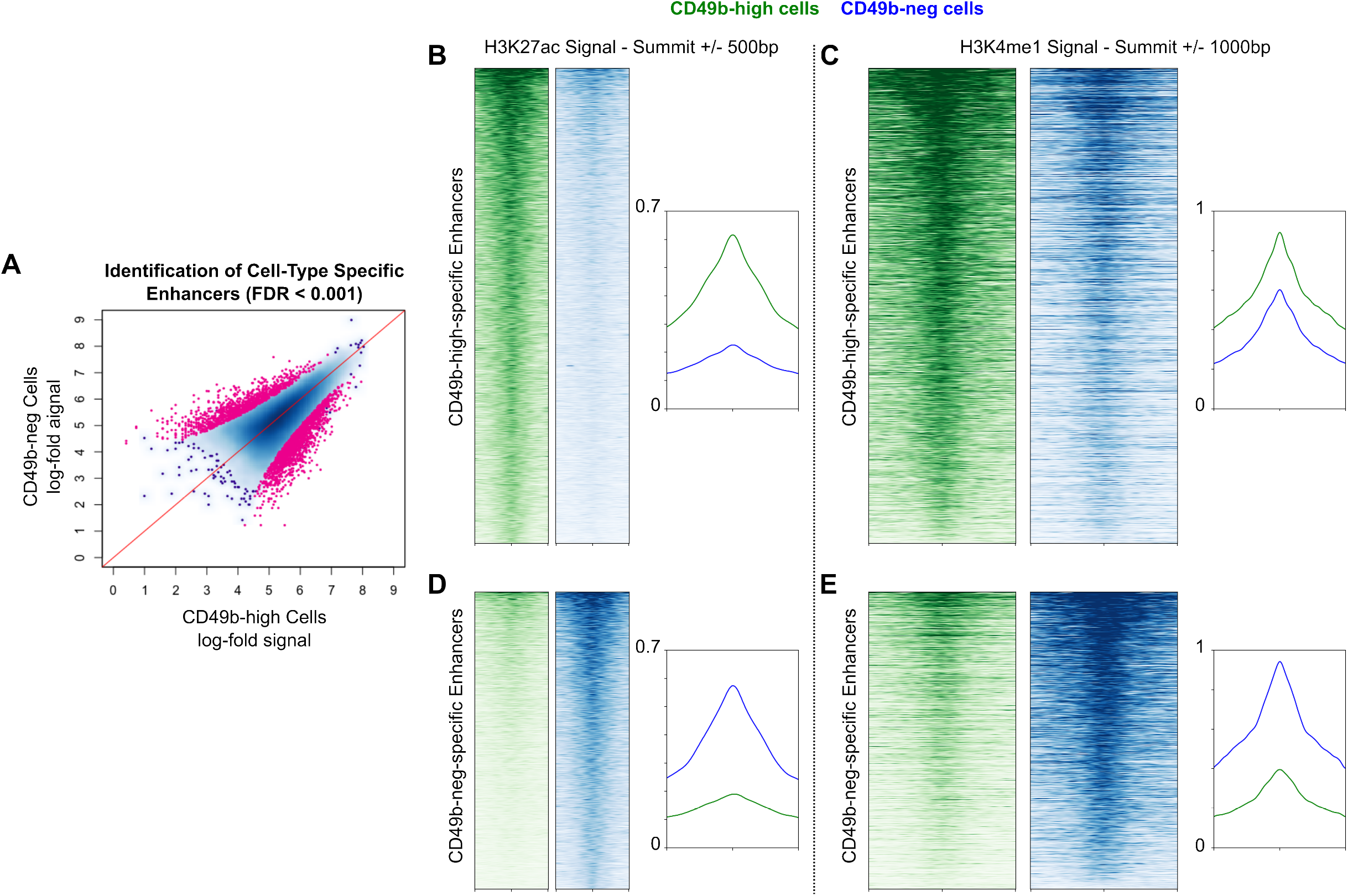
CD49b-neg cells have a distinct repertoire of active enhancers, but maintain the priming H3K4me1 mark on enhancers active in CD49b-high cells. A) Identification of enhancers with greater H3K27ac signal in CD49b-neg or CD49-high cells. Greater distance below the red line indicates greater specificity for CD49b-high cells, and greater distance above the red line indicates greater specificity for CD49b-neg cells. Enhancers with a false discovery rate < 0.001 were deemed specific to one population. B) Heatmap and profile plot showing H3K27ac signal at CD49b-high specific enhancers in CD49b-high (green) and CD49b-neg (blue) cells. For B and D, Heatmaps are centered at enhancer summits and show 500 base pairs up- and downstream. C) Heatmap and profile plot showing H3K4me1 signal at CD49b-high specific enhancers in CD49b-high (green) and CD49b-neg (blue) cells. For C and E, Heatmaps are centered at enhancer summits and show 1000 base pairs up- and downstream. D) Heatmap and profile plot showing H3K27ac signal at CD49b-neg specific enhancers in CD49b-high (green) and CD49b-neg (blue) cells. E) Heatmap and profile plot showing H3K4me1 signal at CD49b-neg specific enhancers in CD49b-high (green) and CD49b-neg (blue) cells.

However, when we examined signal of the priming mark H3K4me1 at the same loci, we found that CD49-neg cells maintain this mark, albeit to a slightly diminished extent, at enhancers active in CD49b-high cells (Fig. 4C). Contrary to our expectation, CD49b-neg cells thus maintain CD49b-high-specific enhancers in a poised state. A similar trend was observed at CD49b-neg enhancers, although to a lesser degree (Fig. 4E). This implies that CD49b-neg neuroblastoma cells, despite having a gene expression program characteristic of differentiated neurons, remain primed to de-differentiate to an immature state.

### CD49b-neg N2a cells maintain H3K4me1 markers at super enhancers that define the CD49b-high state

SEs are large regions of chromatin that often overlap genes controlling cell identity. We used the ROSE algorithm (Whyte et al., 2013) to identify all SEs active in N2a cells, and then used DEseq2 with a stringent FDR (<10e-6) to identify SEs active in the CD49b-neg and CD49b-high populations. This approach identified 69 SEs that are active only in CD49b-neg cells, and 228 SEs active only in CD49b-high cells (Fig. 5A). Profile plots reveal that H3K27ac signal is diminished in each population at SE loci that are active in the opposite cell type (Fig. 5B,C). This is also seen for H3K4me1 signal in SEs active in CD49b-neg cells (Fig. 5E). However, in CD49b-neg cells, H3K4me1 signal at CD49b-high SE loci was only slightly diminished (Fig. 5D). This is illustrated by the SE region overlapping the *Itga2* transcriptional start site, which is one of the SEs that defines the CD49b-high cell state (Fig. 5F). Taken together with data in Fig. 4, these results indicate that CD49b-neg neuroblastoma cells are epigenetically poised to activate the enhancers and SEs that determine the CD49b-high state.

**Figure 5.**
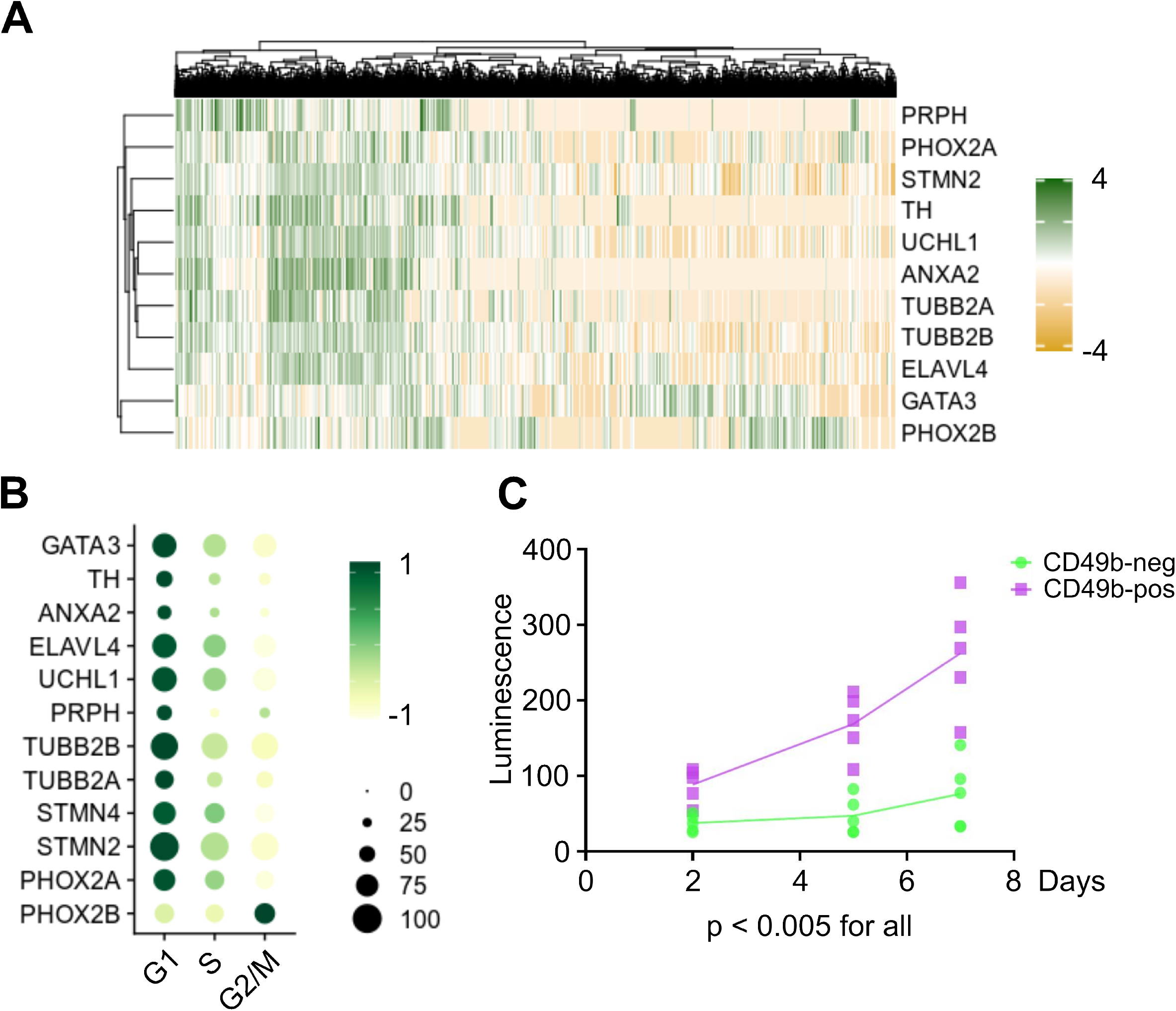
CD49b-negative cells have a distinct super enhancer profile, but maintain H3K4me1 marks on within super enhancers active in CD49b-high cells. A) Identification of super enhancers with greater H3K27ac signal in CD49b-neg or CD49-high cells. Greater distance below the red line indicates greater specificity for CD49b-high cells, and greater distance above the red line indicates greater specificity for CD49b-neg cells. Super enhancers with a false discovery rate < 0.000001 were deemed specific to one population. B) Profile plot showing H3K27ac signal at CD49b-high specific super enhancers in CD49b-high (green) and CD49b-neg (blue) cells. For B-E, Super enhancers are scaled to 25,000 base pairs, and 12,500 base pairs up- and downstream are shown. C) Profile plot showing H3K27ac signal at CD49b-neg specific super enhancers in CD49b-high (green) and CD49b-neg (blue) cells. D) Profile plot showing H3K4me1 signal at CD49b-high specific super enhancers in CD49b-high (green) and CD49b-neg (blue) cells. E) Profile plot showing H3K4me1 signal at CD49b-neg specific super enhancers in CD49b-high (green) and CD49b-neg (blue) cells. F) Tracks plot showing signal for H3K27ac, H3K4me1, and input at the *Itga2* super enhancer in N2a cells. CD49b-high cells are green, CD49b-neg cells are blue. Solid blue bar at the bottom indicates the super enhancer region, and hashed blue line indicates *Itga2* coding region. Shaded grey regions are hand-selected to highlight diminished H3K27ac signal in CD49b-neg cells despite maintenance of H3K4me1 signal. All windows are scaled equally.

### *Neuroblastoma cells can interconvert between CD49b expression states* in vitro

Given the persistence of H3K4me1 marks at CD49b-high-specific enhancers in CD49b-neg N2a cells, we hypothesized that these cells can switch to a CD49b-high state. To test this, we sorted N2a cells and SH-SY5Y cells using the gating strategies shown in Fig. S1, then returned the sorted populations to culture under normal growth conditions for 7 days (N2a cells) or 21 days (SH-SY5Y cells). Cells were then reanalyzed for CD49b expression by flow cytometry. Because N2a cells show a continuum of CD49b expression, we included cells that were neither CD49b-neg nor CD49b-high in our analysis, and referred to this middle population as CD49b-low (Fig. S1A). Consistent with our hypothesis, cultures beginning with a pure population of CD49b-neg cells generate large numbers of cells expressing the antigen in both N2a cells (Fig. 6A,B) and SH-SY5Y cells (Fig. 6C,D). Interestingly, enhancer priming via H3K4me1 may not be necessary for plasticity, as cultures of CD49b-high N2a cells, which do not have strong H3K4me1 signal at CD49b-neg enhancers (Fig. 4E), also give rise to cells lacking the antigen (Fig. 6A-D), suggesting the CD49b-high to CD49b-neg switch involves *de novo* enhancer selection.

**Figure 6.**
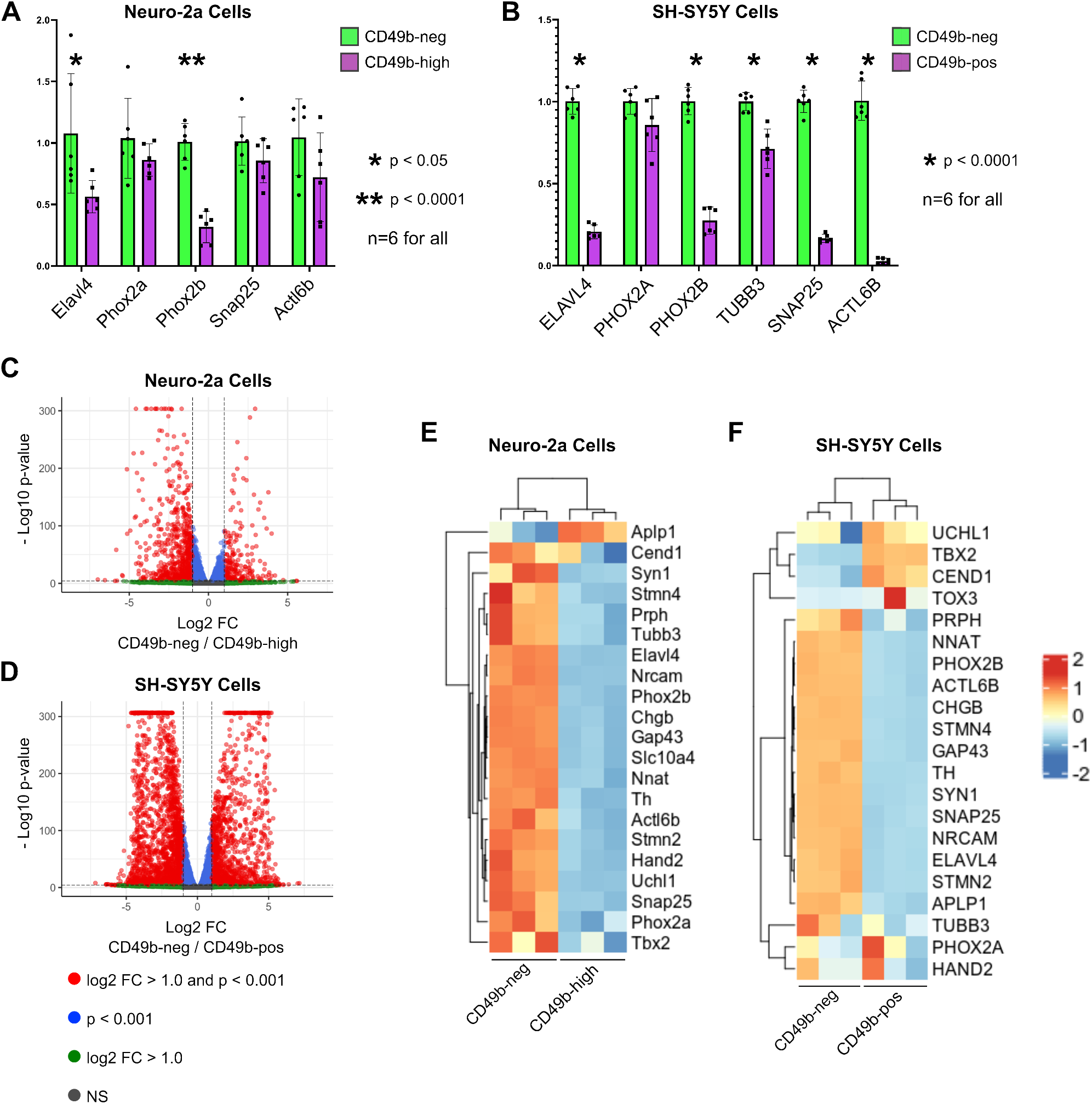
Neuroblastoma cells transition between CD49b expression profiles in culture. A) Representative flow cytometry plots showing reanalysis of N2a cells 7 days after cultures were initiated with the indicated sorted population. B) Quantification of the proportion of N2a cells in each CD49b expression category 7 days after cultures were initiated with the indicated sorted population. Bars indicate mean, dots indicate individual replicates, and error bars indicate standard deviation. C) Representative flow cytometry plots showing reanalysis of SH-SY5Y cells 21 days after cultures were initiated with the indicated sorted population. E) Quantification of the proportion of SH-SY5Y cells in each CD49b expression category 21 days after cultures were initiated with the indicated sorted population. Bars indicate mean, dots indicate individual replicates, and error bars indicate standard deviation.

### *CD49b-neg cells switch to a CD49b-high phenotype and generate tumors* in vivo

We next sought to determine whether neuroblastoma cells exhibit similar phenotypic plasticity *in vivo*. The N2a cell line is derived from a spontaneous tumor in A/J mice, and when injected into syngenic animals forms rapidly-growing tumors in a native microenvironment (Lee et al., 2012; Srinivasan et al., 2018). Taking advantage of this model, we sorted N2a cells into CD49b-neg and CD49b-high populations and then injected 2 × 10^5^ cells per animal into the flank of A/J mice. Mice were euthanized ten days later. Surprisingly, there was no statistically significant difference in weight between tumors grown from CD49b-neg and CD49b-pos cells, although there was greater variability in tumor size in the CD49b-neg group (Fig. 7A). Histological examination of the tumors revealed a lack of neuropil in all cases, suggesting poorly differentiated neuroblastomas, although eosinophilic cytoplasm does appear more abundant in CD49b-high tumors (Fig. 7B,C). Interestingly, both sets of tumors were diffusely positive for CD49b (Fig. 7B,C), confirming that CD49b-neg cells can generate tumors with a CD49b-high phenotype *in vivo*. Both sets of tumors were also diffusely positive for Ki-67 (Fig. 7B,C), indicating significant cell proliferation, although Ki-67 staining was diminished in areas of necrosis (Fig. 7C).

**Figure 7.**
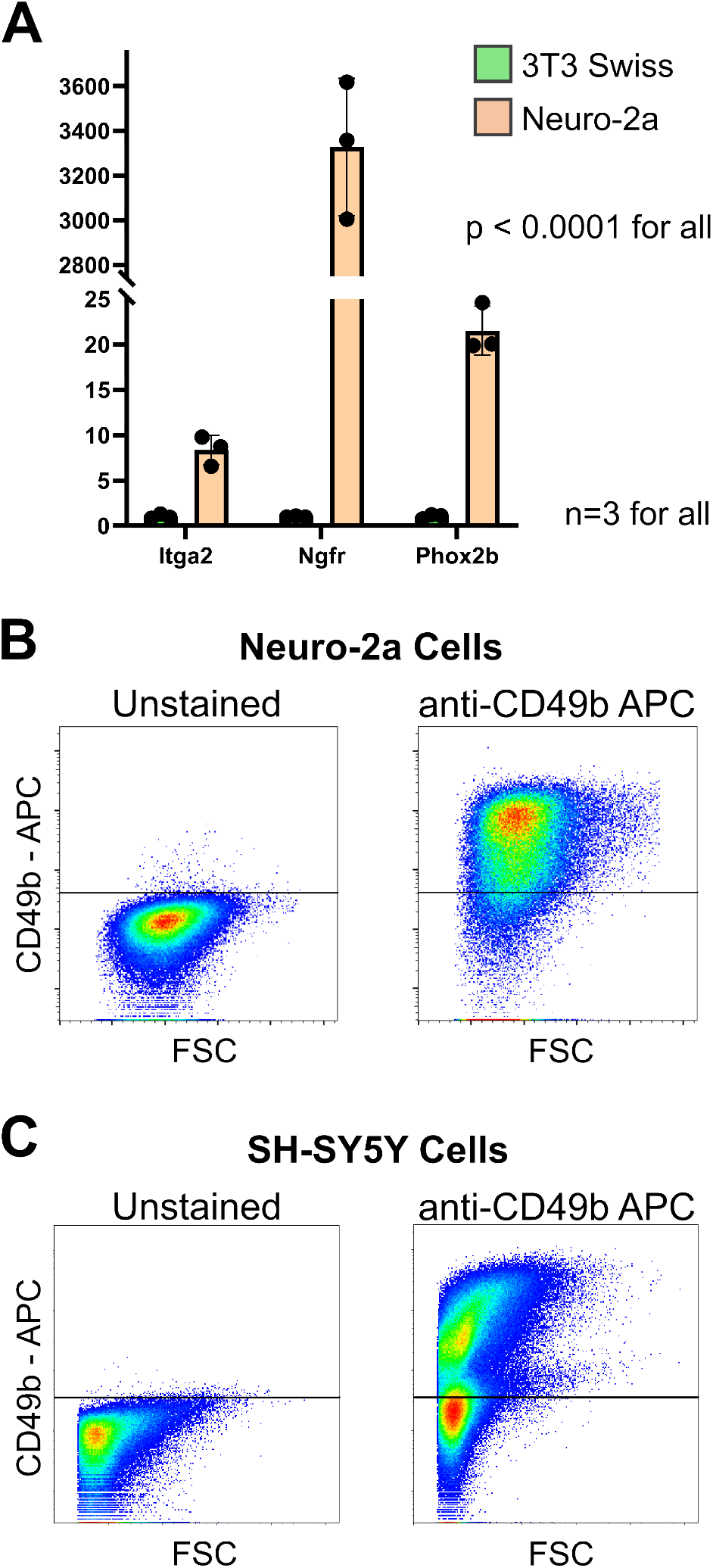
CD49b-neg cells and CD49b-high cells both form CD49b-expressing tumors *in vivo*. A) Tumor weight immediately after euthanizing mice. Bars indicate mean, dots indicate individual replicates, and error bars indicate standard deviation. B) Microscopic examination of tumors grown from CD49b-neg cells. Immunostaining for CD49b (bottom left) and Ki-67 (bottom right) shows that tumors are diffusely reactive for CD49b and highly proliferative. C) Microscopic examination of tumors grown from CD49b-high cells. High-power images of H&E stains show that there is more eosinophilic cytoplasm that in tumors grown from CD49b-neg cells (top right). Immunostaining for CD49b (bottom left) and Ki-67 (bottom right) shows that tumors are diffusely reactive for CD49b and highly proliferative, except in areas of necrosis.

## Discussion

Recent studies have shown that neuroblastoma cells have two predominant states, a differentiated adrenergic state characterized by neuronal transcripts and a less-differentiated mesenchymal state expressing genes seen in neural crest cells. Prior studies assessing these states in neuroblastoma cell lines have sequenced bulk, unsorted populations, making it impossible to investigate heterogeneity within populations (Gartlgruber et al., 2021; van Groningen et al., 2017). Our work establishes that two neuroblastoma cell lines lacking *MYCN* amplification, murine N2a and human SH-SY5Y, contain cells with gene expression patterns characteristic of both the adrenergic and mesenchymal states. We have also shown that N2a and SH-SY5Y cells are plastic between these states. It is known that treating neuroblastoma cells with cytotoxic agents can alter gene expression (Boeva et al., 2017), but prior research in this area has relied on bulk expression profiles, so until now it has not been clear whether individual cells are plastic. We have now shown that sorted populations of neuroblastoma cells revert to a mixed population, meaning that cells either directly switch phenotypes or undergo asymmetric divisions producing disparate daughter cells. Reversion towards equilibrium proportions from sorted starting populations has been shown previously for cancer cells (Gupta et al., 2011), but to our knowledge this is the first demonstration in neuroblastoma.

We have shown that the adrenergic and mesenchymal states can be distinguished by expression of the CD49b antigen. CD49b is an integrin that marks immature neuronal precursors in the peripheral nervous system (Belkind-Gerson et al., 2015; Joseph et al., 2011). Downregulation of this antigen is associated with a transition to a committed neuronal state, but reversion of CD49b-negative neurons to a CD49b-expressing state has not been observed during normal development. The ability of tumor cells to revert from a CD49b-neg state, which is characterized by expression of neuronal genes, inverts the normal trajectory of neurogenesis. We found that CD49b-neg N2a cells maintain the priming H3K4me1 mark at enhancer loci that are active in CD49b-high cells, which may explain their ability to revert to a less mature gene expression state. Decommissioning of primed enhancers appears to be a key mechanism for cells to maintain differentiation (Tao et al., 2021; Whyte et al., 2012). We thus speculate that oncogenic mutations in neuroblastoma likely act, at least in part, by preventing proper decommissioning of enhancer elements. The recent report that loss of the tumor suppressor gene *ARID1A* causes de-differentiation of neuroblastoma cells supports this hypothesis (Shi et al., 2020).

SEs are large genomic regions marked heavily by H3K27ac and densely bound by transcription factors (Parker et al., 2013; Whyte et al., 2013). These regions control cell fate-determining genes, and disruptions to SEs occur in cancers including neuroblastoma (Gartlgruber et al., 2021; Lovén et al., 2013; van Groningen et al., 2017). We found that neuroblastoma cells marked by high or low CD49b expression have distinct SE profiles, which is strong evidence that the CD49b antigen distinguishes different biological states. Additionally, one of the SEs we identified in CD49b-high cells overlaps with the *Itga2* gene, which encodes the CD49b antigen. This suggests that CD49b is not merely a marker of proliferative, immature cells, but may have an important role in establishing such a state. Intriguingly, CD49b activates the AKT pathway in esophageal squamous cell carcinoma and hepatocellular carcinoma (Huang et al., 2021; Juratli et al., 2022), suggesting a mechanistic link between the antigen and a correlated signaling network. While functional studies of CD49b are outside the scope of this report, we anticipate it will be an interesting avenue for future studies.

Although SEs active in CD49b-high cells have diminished H3K27ac signal in CD49b-low cells, we found these same SEs to maintain high levels of H3K4me1. Similarly, maintenance of the H3K4me2 priming mark at SEs has been observed across macrophage subtypes (Gosselin et al., 2014). H3K4me1 marks act by recruiting the BAF complex to ordinary enhancers (Local et al., 2018), where the BAF complex establishes accessible chromatin as a key step in enhancer activation (Vierbuchen et al., 2017). To our knowledge, whether the H3K4me1 mark plays a priming role in SEs as well as at traditional enhancers has not been explored. We speculate that multipotent cells may maintain alternate fate potentials by mono-or dimethylating H3K4 in SE regions, thus recruiting the BAF complex and generating open chromatin that can be rapidly activated to initiate expression of novel fate-determining transcripts. This model would correlate with the recent description of domains of regulatory chromatin (DORCs), which are large regions around fate-determining genes with an open chromatin structure (Ma et al., 2020). DORCs correlate highly with SEs identified by H3K27ac signal, and chromatin accessibility at DORCs generally precedes gene expression. Future work to test these hypotheses and establish the underlying mechanisms could yield valuable insights for neuroblastoma therapy.

## Materials and Methods

### Cell Lines and Cell Culture

3T3 Swiss (CCL-92), N2a (CCL-131) and SH-SY5Y (CRL-2266) cells without mycoplasma detection were obtained from ATCC (Manassas, VA). SH-SY5Y cell identity was confirmed by short tandem repeat testing by ATCC. 3T3 cells were maintained in DMEM media (#11965-118 ThermoFisher, Waltham, MA) supplemented with 10% fetal bovine serum (FBS) and 1% penicillin-streptomycin (pen-strep). N2a cells were maintained in EMEM media (#12-611F Lonza, Basil, Switzerland) supplemented with 10% FBS and 1% pen-strep. SH-SY5Y cells were maintained in DMEM/F12 (1:1) media (#11330-032 ThermoFisher) supplemented with 15% FBS and 1% pen-strep. Cells were monitored daily and passaged when they reached 80% confluency. All experiments were performed on cells less than ten passages from delivery from ATCC.

### Flow Cytometry and Cell Sorting

All flow cytometry and cell sorting was performed at the Harvard Stem Cell Institute Center for Regenerative Medicine Flow Cytometry Core facility on FACSAria instruments (BD Biosciences, East Rutherford, NJ). Cells were trypsinized, spun down, resuspended in PBS plus 10% FBS, and counted. Cells were blocked in PBS with 10% FBS on ice for 20 minutes, stained for 20 minutes on ice with the following antibodies at the following concentrations: anti-mouse CD49b 1:500 (#103511 Biolegend, San Diego, CA), anti-human CD49b 1:750 (#359310 Biolegend), anti-mouse gp130 1:100 (#149404 Biolegend), anti-human gp130 1:100 (#362010 Biolegend), anti-phospho-Akt 1:50 (#4071 Cell Signaling Technology, Danvers, MA). Scatter profile and DAPI staining were used to exclude debris, doublets, and dead cells. Gating for analysis was performed as described in the Results section. All analysis was conducted using FlowJo software (BD Biosciences).

### Quantitative PCR

Total RNA was isolated from cells using RNEasy mini kits (Qiagen, Germantown, MD). Reverse transcription and cDNA amplification were done using the iTaq Universal SYBR Green One-Step Kit (Bio-Rad, Hercules, CA) and a CFX96 real-time system (Bio-Rad, Hercules, CA). Primer sequences for the murine genes *Itga2, Ngfr, Elavl4, Phox2a, Phox2b, Snap25, Actl6b*, and *Gapdh* (normalization control) and for the human genes *ELAVL4, PHOX2A, PHOX2B, SNAP25, ACTL6B, TUBB3*, and *GAPDH* (normalization control) were obtained from the Harvard Medical School PCR PrimerBank. Primer sequences are included in Supplementary Table 1.

### RNA Sequencing and Data Analysis

Cells were isolated by sorting, after which total RNA was isolated using RNEasy mini kits (Qiagen) and treated with RNase-free DNase (Qiagen). RNA was quantified using the Qubit RNA HS Assay Kit (ThermoFisher). Poly(A)-enriched libraries for sequencing were then generated using the NEBNext Ultra II Directional RNA Library Prep Kit for Illumina and the NEBNext Poly(A) mRNA Magnetic Isolation Module (New England Biolabs, Ipswich, MA). Paired-end sequencing was performed on the Illumina NextSeq instrument at the Harvard University Bauer Core. Reads were aligned to the mm10 (mouse) and hg38 (human) reference genomes using STAR 2.7.3 on the Mass General Brigham ERISOne Cluster. Counts were computed using featureCounts 2.0.3 on the ERISOne Cluster.

Differential gene expression analysis was performed using the DESeq2 package in R 4.1.0, with count normalization done by DESeq2’s default median of ratios method. Heatmaps were generated with the ComplexHeatmap R package, and volcano plots were generated with the EnhancedVolcano R package. Gene Set Enrichment Analysis (GSEA) was performed using the clusterProfiler R package.

### CUT&RUN and Data Analysis

Cells were isolated by sorting, after which the CUT&RUN assay was performed using the CUT&RUN Assay Kit (Cell Signaling Technology). The following antibodies were used at 1:50 dilution: H3K4me1 (#5326, Cell Signaling Technology) and H3K27ac (#8173, Cell Signaling Technology). Sequencing libraries were built using DNA Library Prep Kit for Illumina (Cell Signaling Technology, Danvers, MA). Paired-end sequencing was performed on the Illumina NextSeq instrument at the Harvard University Bauer Core. Reads were aligned to the mm10 (mouse) and hg38 (human) reference genomes using Bowtie 2.4.1 on the Mass General Brigham ERISOne Cluster. SAM files output by Bowtie were converted to BAM format using Samtools View in the Samtools 1.4.1 package, and BAM files were filtered for uniquely mapped reads using Sambamba 0.4.7. BAM files were then randomly downsampled to contain equivalent numbers of reads using Samtools View. Peak calling was performed on downsampled BAM files using the MACS2 algorithm version 2.1. H3K27ac peaks with differential signal in CD49b-neg and CD49b-high cells were then identified using the DiffBind package in R 4.1.0. Super enhancers (SEs) were identified using the ROSE algorithm on downsampled H3K27ac BAM files (Whyte et al., 2013) in Python 2.7.3, and SEs specifically active in CD49b-neg and CD49-high cells were then identified using DiffBind. Heatmaps and profile plots were generated using deepTools 3.5.1. Genome browser tracks were created with the Integrative Genomics Viewer 2.11.1 (Robinson et al., 2011).

### Published scRNA-seq Data

Previously-published scRNA-seq data of primary neuroblastoma lesions was queried (Dong et al., 2020). Data are available from the NCBI Gene Expression Omnibus under accession GSE137804. Raw data was obtained from the NCBI Sequence Run Archive using the SRA Toolkit “fastq-dump” command. Genome alignment and feature-barcode matrix generation was performed with the Cell Ranger “cellranger count” command on the Mass General Brigham ERISOne Cluster. Further analysis was performed with Seurat in the R environment. Patient samples T10, T34, T69, T71, and T92 were selected for being high-risk lesions lacking *MYCN* amplification. Tumor cells were identified based on cellular barcodes provided in meta data from the depositing authors. Single-cell heatmaps were generated using the ComplexHeatmap R package, and dot plots were generated using the Seurat package’s DotPlot function.

### Cell Proliferation Assay

5×10^3^ SH-SY5Y cells were plated in each well of 48-well culture dishes after sorting into CD49b-neg and CD49b-pos populations. At the indicated time points, cell viability was assayed using the CellTiter-Glo Luminescent Cell Viability Assay (Promega, Madison, WI).

### In Vivo Tumor Model

All animal procedures were approved by the Institutional Animal Care and Use Committee at Massachusetts General Hospital. A/J albino mice (strain #000646, Jackson Laboratory, Bar Harbor, ME) at 3 weeks of age were injected on the right flank with 10^5^ N2a cells sorted into either CD49b-neg or CD49b-high populations. 5 mice were injected with each group of cells. Ten days after injection, mice were euthanized and tumors were explanted. Tumors were weighed and fixed overnight in 10% formalin, after which they were transferred to 70% ethanol for long-term storage.

### Tumor Histology

Fixed tumors were sectioned, mounted on slides, and stained with hematoxylin and eosin or with antibodies against Ki-67 or CD49b by the staff at the Histopathology Research Core at Massachusetts General Hospital. Slides were reviewed and imaged by a board-certified pathologist with expertise in neuroblastoma (KMC).

### Statistical Analysis

Statistical comparisons were performed using the unpaired, two-tailed t-test with p<0.05 set as the predetermined level of significance. Statistical testing was performed using GraphPad Prism (San Diego, CA).

## Acknowledgments

The authors thank the staff at the Harvard University Bauer Core, Harvard Stem Cell Institute Center for Regenerative Medicine Flow Cytometry Core, and the MGH Histopathology Research Core for their expertise and assistance.

## Data Availability

The datasets generated for this study can be found in the NCBI GEO under accession NUMBER.

## Author Contributions

Conceptualization, RAG and AMG; Methodology, RAG, NP, and JLM; Formal Analysis, RAG, NP, JLM, and KMC; Investigation, RAG, NP, and JLM; Resources, AMG and RH; Data curation, RAG and NP; Writing – original draft, RAG and AMG; Writing – review and editing, RAG, AMG, NP, JLM, and AJM; Visualization, RAG and NP; Supervision, AMG and RH; Project Administration, AMG; Funding Acquisition, RAG and AMG.

**Supplemental Figure 1. Gating strategies for identifying CD49b populations**

A) Representative examples of unstained (left) and CD49b-stained (right) N2a cells, with the gates used to identify CD49b-neg, CD49b-low, and CD49b-high cells.

B) Representative examples of unstained (left) and CD49b-stained (right) SH-SY5Y cells, with the gates used to identify CD49b-neg and CD49b-pos cells.

**Supplemental Figure 2. Expression of transcription factor genes associated with adrenergic and mesenchymal neuroblastoma cell states, as described by van Groningen et al (2017)**

A) Heatmaps showing expression of genes encoding for transcription factors associated with the adrenergic neuroblastoma cell state, showing enrichment in CD49b-neg cells in the indicated cell lines.

B) Heatmaps showing expression of genes encoding for transcription factors associated with the mesenchymal neuroblastoma cell state, showing enrichment in CD49b-pos/high cells in the indicated cell lines.

**Supplemental Figure 3. CD49b differentiates neuroblastoma cells with different signaling milieus**

A)GSEA enrichment plots showing enrichment of genes associated with the P I3K-AKT Signaling Pathway GSEA term CD49b-high or CD49b-pos cells in the indicate cell lines relative to CD49b-neg cells.

B) GSEA enrichment plots showing enrichment of genes associated with the Cytokine-Cytokine Receptor Interaction GSEA term in CD49b-high or CD49b-pos cells the indicated cell lines relative to CD49b-neg cells.

C) Heatmaps showing enrichment of genes associated with PI3K-AKT Signaling Pathway GSEA term in CD49b-pos or CD49b-high populations of the indicate cell lines.

D) Heatmaps showing enrichment of genes associated with Cytokine-Cytokine Receptor Interaction GSEA term in CD49b-pos or CD49b-high populations of the indicate cell lines.

E) Quantification of flow cytometry analysis demonstrating increased p-Akt/p-AKT levels in CD49b-expresing cells relative to CD49b-neg cells. Bars indicate mean, dots indicate individual replicates, and error bars indicate standard deviation.

F) Quantification of flow cytometry analysis demonstrating increased gp130 levels in CD49b-expresing cells relative to CD49b-neg cells. Bars indicate mean, dots indicate individual replicates, and error bars indicate standard deviation.

**Supplemental Table 1. qPCR primer sequences**

**Supplemental Table 2. GSEA pathway enrichment in N2a CD49b-neg cells relative to CD49b-high cells**

**Supplemental Table 3. GSEA pathway enrichment in SH-SY5Y CD49b-neg cells relative to CD49b-pos cells**

## References

Abe, S., Yamaguchi, S., Sato, Y. and Harada, K. (2016). Sphere-Derived Multipotent Progenitor Cells Obtained From Human Oral Mucosa Are Enriched in Neural Crest Cells. Stem Cells Transl Med 5, 117–128.

Baker, D. L., Reddy, U. R., Pleasure, D., Thorpe, C. L., Evans, A. E., Cohen, P. S. and Ross, A. H. (1989). Analysis of nerve growth factor receptor expression in human neuroblastoma and neuroepithelioma cell lines. Cancer Res 49, 4142–4146.

Belkind-Gerson, J., Hotta, R., Nagy, N., Thomas, A. R., Graham, H., Cheng, L., Solorzano, J., Nguyen, D., Kamionek, M., Dietrich, J., et al. (2015). Colitis Induces Enteric Neurogenesis Through a 5-HT4–dependent Mechanism: Inflammatory Bowel Diseases 21, 870–878.

Boeva, V., Louis-Brennetot, C., Peltier, A., Durand, S., Pierre-Eugène, C., Raynal, V., Etchevers, H. C., Thomas, S., Lermine, A., Daudigeos-Dubus, E., et al. (2017). Heterogeneity of neuroblastoma cell identity defined by transcriptional circuitries. Nature Genetics 49, 1408– 1413.

Cohn, S. L., Pearson, A. D. J., London, W. B., Monclair, T., Ambros, P. F., Brodeur, G. M., Faldum, A., Hero, B., Iehara, T., Machin, D., et al. (2016). The International Neuroblastoma Risk Group (INRG) Classification System: An INRG Task Force Report. Journal of Clinical Oncology.

Cotterman, R. and Knoepfler, P. S. (2009). N-Myc Regulates Expression of Pluripotency Genes in Neuroblastoma Including lif, klf2, klf4, and lin28b. PLOS ONE 4, e5799.

Creyghton, M. P., Cheng, A. W., Welstead, G. G., Kooistra, T., Carey, B. W., Steine, E. J., Hanna, J., Lodato, M. A., Frampton, G. M., Sharp, P. A., et al. (2010). Histone H3K27ac separates active from poised enhancers and predicts developmental state. PNAS 107, 21931–21936.

Crispatzu, G., Rehimi, R., Pachano, T., Bleckwehl, T., Cruz-Molina, S., Xiao, C., Mahabir, E., Bazzi, H. and Rada-Iglesias, A. (2021). The chromatin, topological and regulatory properties of pluripotency-associated poised enhancers are conserved in vivo. Nat Commun 12, 4344.

Dong, R., Yang, R., Zhan, Y., Lai, H.-D., Ye, C.-J., Yao, X.-Y., Luo, W.-Q., Cheng, X.-M., Miao, J.-J., Wang, J.-F., et al. (2020). Single-Cell Characterization of Malignant Phenotypes and Developmental Trajectories of Adrenal Neuroblastoma. Cancer Cell 38, 716-733.e6.

Finklestein, J. Z., Klemperer, M. R., Evans, A., Bernstein, I., Leikin, S., McCreadie, S., Grosfeld, J., Hittle, R., Weiner, J., Sather, H., et al. (1979). Multiagent chemotherapy for children with metastatic neuroblastoma: A report from childrens cancer study group. Medical and Pediatric Oncology 6, 179–188.

Gartlgruber, M., Sharma, A. K., Quintero, A., Dreidax, D., Jansky, S., Park, Y.-G., Kreth, S., Meder, J., Doncevic, D., Saary, P., et al. (2021). Super enhancers define regulatory subtypes and cell identity in neuroblastoma. Nature Cancer 2, 114–128.

Gosselin, D., Link, V. M., Romanoski, C. E., Fonseca, G. J., Eichenfield, D. Z., Spann, N. J., Stender, J. D., Chun, H. B., Garner, H., Geissmann, F., et al. (2014). Environment Drives Selection and Function of Enhancers Controlling Tissue-Specific Macrophage Identities. Cell 159, 1327–1340.

Grosfeld, J. L., Schatzlein, M., Ballantine, T. V. N., Weetman, R. M. and Baehner, R. L. (1978). Metastatic neuroblastoma: Factors influencing survival. Journal of Pediatric Surgery 13, 59–65.

Gupta, P. B., Fillmore, C. M., Jiang, G., Shapira, S. D., Tao, K., Kuperwasser, C. and Lander, E. S. (2011). Stochastic State Transitions Give Rise to Phenotypic Equilibrium in Populations of Cancer Cells. Cell 146, 633–644.

Hatzi, E., Murphy, C., Zoephel, A., Ahorn, H., Tontsch, U., Bamberger, A.-M., Yamauchi-Takihara, K., Schweigerer, L. and Fotsis, T. (2002). N-myc oncogene overexpression down-regulates leukemia inhibitory factor in neuroblastoma. European Journal of Biochemistry 269, 3732–3741.

Helmsauer, K., Valieva, M. E., Ali, S., Chamorro González, R., Schöpflin, R., Röefzaad, C., Bei, Y., Dorado Garcia, H., Rodriguez-Fos, E., Puiggròs, M., et al. (2020). Enhancer hijacking determines extrachromosomal circular MYCN amplicon architecture in neuroblastoma. Nat Commun 11, 5823.

Huang, W., Zhu, J., Shi, H., Wu, Q. and Zhang, C. (2021). ITGA2 Overexpression Promotes Esophageal Squamous Cell Carcinoma Aggression via FAK/AKT Signaling Pathway. OTT 14, 3583–3596.

Joseph, N. M., He, S., Quintana, E., Kim, Y.-G., Núñez, G. and Morrison, S. J. (2011). Enteric glia are multipotent in culture but primarily form glia in the adult rodent gut. J. Clin. Invest. 121, 3398–3411.

Juratli, M. A., Zhou, H., Oppermann, E., Bechstein, W. O., Pascher, A., Chun, F. K.-H., Juengel, E., Rutz, J. and Blaheta, R. A. (2022). Integrin α2 and β1 Cross-Communication with mTOR/ AKT and the CDK-Cyclin Axis in Hepatocellular Carcinoma Cells. Cancers 14, 2430.

Kildisiute, G., Kholosy, W. M., Young, M. D., Roberts, K., Elmentaite, R., Hooff, S. R. van, Pacyna, C. N., Khabirova, E., Piapi, A., Thevanesan, C., et al. (2021). Tumor to normal single-cell mRNA comparisons reveal a pan-neuroblastoma cancer cell. Science Advances 7, eabd3311.

Lee, P., Zhang, R., Li, V., Liu, X., Sun, R. W., Che, C.-M. and Wong, K. K. (2012). Enhancement of anticancer efficacy using modified lipophilic nanoparticle drug encapsulation. Int J Nanomedicine 7, 731–737.

Lee, J. W., Son, M. H., Cho, H. W., Ma, Y. E., Yoo, K. H., Sung, K. W. and Koo, H. H. (2018). Clinical significance of MYCN amplification in patients with high-risk neuroblastoma. Pediatr Blood Cancer 65, e27257.

Local, A., Huang, H., Albuquerque, C. P., Singh, N., Lee, A. Y., Wang, W., Wang, C., Hsia, J. E., Shiau, A. K., Ge, K., et al. (2018). Identification of H3K4me1-associated proteins at mammalian enhancers. Nat Genet 50, 73–82.

Long, H. K., Prescott, S. L. and Wysocka, J. (2016). Ever-Changing Landscapes: Transcriptional Enhancers in Development and Evolution. Cell 167, 1170–1187.

Lovén, J., Hoke, H. A., Lin, C. Y., Lau, A., Orlando, D. A., Vakoc, C. R., Bradner, J. E., Lee, T. I. and Young, R. A. (2013). Selective Inhibition of Tumor Oncogenes by Disruption of Super-Enhancers. Cell 153, 320–334.

Ma, S., Zhang, B., LaFave, L. M., Earl, A. S., Chiang, Z., Hu, Y., Ding, J., Brack, A., Kartha, V. K., Tay, T., et al. (2020). Chromatin Potential Identified by Shared Single-Cell Profiling of RNA and Chromatin. Cell 183, 1103-1116.e20.

Morarach, K., Mikhailova, A., Knoflach, V., Memic, F., Kumar, R., Li, W., Ernfors, P. and Marklund, U. (2021). Diversification of molecularly defined myenteric neuron classes revealed by single-cell RNA sequencing. Nature Neuroscience 24, 34–46.

Newman, E. A., Abdessalam, S., Aldrink, J. H., Austin, M., Heaton, T. E., Bruny, J., Ehrlich, P., Dasgupta, R., Baertschiger, R. M., Lautz, T. B., et al. (2019). Update on neuroblastoma. Journal of Pediatric Surgery 54, 383–389.

Okabe, A. and Kaneda, A. (2021). Transcriptional dysregulation by aberrant enhancer activation and rewiring in cancer. Cancer Science 112, 2081–2088.

Parker, S. C. J., Stitzel, M. L., Taylor, D. L., Orozco, J. M., Erdos, M. R., Akiyama, J. A., van Bueren, K. L., Chines, P. S., Narisu, N., NISC Comparative Sequencing Program, et al. (2013). Chromatin stretch enhancer states drive cell-specific gene regulation and harbor human disease risk variants. Proceedings of the National Academy of Sciences 110, 17921–17926.

Paul, P., Volny, N. S., Lee, S., Qiao, J. and Chung, D. H. (2013). Gli1 Transcriptional Activity is Negatively Regulated by AKT2 in Neuroblastoma. Oncotarget 4, 1149–1157.

Rada-Iglesias, A., Bajpai, R., Swigut, T., Brugmann, S. A., Flynn, R. A. and Wysocka, J. (2011). A unique chromatin signature uncovers early developmental enhancers in humans. Nature 470, 279–283.

Robinson, J. T., Thorvaldsdóttir, H., Winckler, W., Guttman, M., Lander, E. S., Getz, G. and Mesirov, J. P. (2011). Integrative Genomics Viewer. Nat Biotechnol 29, 24–26.

Shi, H., Tao, T., Abraham, B. J., Durbin, A. D., Zimmerman, M. W., Kadoch, C. and Look, A. T. (2020). ARID1A loss in neuroblastoma promotes the adrenergic-to-mesenchymal transition by regulating enhancer-mediated gene expression. Sci Adv 6, eaaz3440.

Srinivasan, P., Wu, X., Basu, M., Rossi, C. and Sandler, A. D. (2018). PD-L1 checkpoint inhibition and anti-CTLA-4 whole tumor cell vaccination counter adaptive immune resistance: A mouse neuroblastoma model that mimics human disease. PLOS Medicine 15, e1002497.

Tao, L., Yu, H. V., Llamas, J., Trecek, T., Wang, X., Stojanova, Z., Groves, A. K. and Segil, N. (2021). Enhancer decommissioning imposes an epigenetic barrier to sensory hair cell regeneration. Developmental Cell 56, 2471-2485.e5.

Tirosh, I., Izar, B., Prakadan, S. M., Wadsworth, M. H., Treacy, D., Trombetta, J. J., Rotem, A., Rodman, C., Lian, C., Murphy, G., et al. (2016). Dissecting the multicellular ecosystem of metastatic melanoma by single-cell RNA-seq. Science 352, 189–196.

van Groningen, T., Koster, J., Valentijn, L. J., Zwijnenburg, D. A., Akogul, N., Hasselt, N. E., Broekmans, M., Haneveld, F., Nowakowska, N. E., Bras, J., et al. (2017). Neuroblastoma is composed of two super-enhancer-associated differentiation states. Nature Genetics 49, 1261– 1266.

van Groningen, T., Akogul, N., Westerhout, E. M., Chan, A., Hasselt, N. E., Zwijnenburg, D. A., Broekmans, M., Stroeken, P., Haneveld, F., Hooijer, G. K. J., et al. (2019). A NOTCH feed-forward loop drives reprogramming from adrenergic to mesenchymal state in neuroblastoma. Nat Commun 10, 1530.

Vierbuchen, T., Ling, E., Cowley, C. J., Couch, C. H., Wang, X., Harmin, D. A., Roberts, C. W. M. and Greenberg, M. E. (2017). AP-1 Transcription Factors and the BAF Complex Mediate Signal-Dependent Enhancer Selection. Molecular Cell 68, 1067-1082.e12.

Whyte, W. A., Bilodeau, S., Orlando, D. A., Hoke, H. A., Frampton, G. M., Foster, C. T., Cowley, S. M. and Young, R. A. (2012). Enhancer decommissioning by LSD1 during embryonic stem cell differentiation. Nature 482, 221–225.

Whyte, W. A., Orlando, D. A., Hnisz, D., Abraham, B. J., Lin, C. Y., Kagey, M. H., Rahl, P. B., Lee, T. I. and Young, R. A. (2013). Master Transcription Factors and Mediator Establish Super-Enhancers at Key Cell Identity Genes. Cell 153, 307–319.

Yanishevski, D., McCarville, M. B., Doubrovin, M., Spiegl, H. R., Zhao, X., Lu, Z., Federico, S. M., Furman, W. L., Murphy, A. J. and Davidoff, A. M. (2020). Impact of MYCN status on response of high-risk neuroblastoma to neoadjuvant chemotherapy. Journal of Pediatric Surgery 55, 130–134.

Yoo, M., Choi, K.-Y., Kim, J., Kim, M., Shim, J., Choi, J.-H., Cho, H.-Y., Oh, J.-P., Kim, H.-S., Kaang, B.-K., et al. (2017). BAF53b, a Neuron-Specific Nucleosome Remodeling Factor, Is Induced after Learning and Facilitates Long-Term Memory Consolidation. J. Neurosci. 37, 3686–3697.

